# Sugar-rich foods exacerbate antibiotic-induced microbiome injury

**DOI:** 10.1101/2024.10.14.617881

**Authors:** Anqi Dai, Peter A. Adintori, Tyler Funnell, William P. Jogia, Teng Fei, Nicholas R. Waters, Madhumitha Rangesa, Annamaria Ballweg, Brianna Gipson, Sandeep Raj, Eiko Hayase, Kate A. Markey, Marina Burgos da Silva, Oriana Miltiadous, Corrado Zuanelli Brambilla, Marissa Lubin Buchan, Tatnisha Peets, Ana Gradissimo, Natalie Smith, Zoe Katsamaikis, Adam Warren, Luigi A. Amoretti, Caichen Duan, Chenzhen Zhang, Fanny Matheis, Alexis P. Sullivan, John B. Slingerland, Annelie G. Clurman, Daniel G. Brereton, Paul A. Giardina, Antonio L.C. Gomes, Abigail J. Johnson, Dan Knights, Robert R. Jenq, Miguel-Angel Perales, Sergio A. Giralt, Jonas Schluter, Marcel R.M. van den Brink, Jonathan U. Peled

**Affiliations:** Department of Immunology, Sloan Kettering Institute, Memorial Sloan Kettering Cancer Center, New York, NY, USA; Food and Nutrition Services; Department of Medicine, Adult Bone Marrow Transplant Service, Memorial Sloan Kettering Cancer Center, New York, NY, USA; Institute for Systems Genetics, Department for Microbiology, NYU Grossman School of Medicine, New York, NY, USA; Perlmutter Cancer Center, NYU Grossman School of Medicine, New York, NY, USA; Department of Epidemiology and Biostatistics, Memorial Sloan Kettering Cancer Center, New York, New York; City of Hope, Duarte, CA, USA; Departments of Genomic Medicine and Stem Cell Transplantation Cellular Therapy, Division of Cancer Medicine, University of Texas MD Anderson Cancer Center, Houston, TX, USA; Translational Science and Therapeutics Division, Fred Hutchinson Cancer Center, Seattle, WA, USA and Division of Medicine, University of Washington, Seattle, WA, USA; Department of Pediatrics, Memorial Sloan Kettering Cancer Center, New York, NY; Department of Medical Biotechnologies, University of Siena, Siena, Italy; Hematology Unit, Department of Oncology, Azienda Ospedaliera Universitaria Senese, Siena, Italy; Food and Nutrition Services, MSKCC; Infectious Disease Service, Department of Medicine, Memorial Sloan Kettering Cancer Center, New York, NY, USA; XBiome, Boston, MA, USA; Division of Epidemiology and Community Health, School of Public Health, University of Minnesota, Minneapolis, MN, USA; Department of Computer Science and Engineering, University of Minnesota, Minneapolis, MN, USA; Biotechnology Institute, University of Minnesota, Saint Paul, MN, USA; Weill Cornell Medical College, New York, NY, USA

## Abstract

Intestinal microbiota composition is implicated in several diseases; understanding the factors that influence it are key to elucidating host-commensal interactions and to designing microbiome-targeted therapies. We quantified how diet influences microbiome dynamics in hospitalized patients. We recorded 9,419 meals consumed by 173 patients undergoing hematopoietic cell transplantation and profiled the microbiome in 1,009 longitudinally collected stool samples from 158 of them. Caloric intake was correlated with fecal microbiota diversity. Bayesian inference revealed associations between intake of sweets or sugars during antibiotic exposure with microbiome disruption, as assessed by low diversity or expansion of the pathobiont *Enterococcus*. We validated this observation experimentally, finding that sucrose exacerbated antibiotic-induced *Enterococcus* expansion in mice. Taken together, our results suggest that avoiding sugar-rich foods during antibiotic treatment may reduce microbiome injury.

**One Sentence Summary:** Analyses of hematopoietic cell transplant patients and of mice uncover links between foods and microbiome disruption.

## Introduction

The intestinal microbiota modulates host metabolism and immunity, and perturbations of microbiome composition are linked with various disease states (*1*, *2*). Although a precise definition of a homeostatic human microbiome composition remains elusive (*3*, *4*), a number of disease states share dysbiotic patterns in microbial community compositions, often characterized by loss of diversity and expansion of facultative anaerobes (*5*). Understanding the factors that influence microbiome injury and dysbiosis is key to deciphering the interactions between host and commensal organisms and to designing therapeutic strategies that target the microbiome.

Host and environmental factors shape intestinal microbiome compositions; in particular, a major effect of diet has been observed both in mice and humans (*6–17*). Most of the human analyses, however, were conducted in volunteers in small dietary-intervention trials in subjects with chronic conditions or healthy volunteers with unperturbed, stable microbiomes (*18*). Severe perturbations, such as those which occur during acute illness or intensive medical treatment (*4*, *19–22*) are less well understood. Also, while animal studies suggest that dietary perturbations exert effects on microbial composition within hours (*6*, *23–25*) most human studies have correlated fecal microbiome compositions with long-term habitual diet (*9*, *10*, *26–29*) or variations on months-long timescales (*30*, *31*) and relied on recall-based surveys whose imprecision and limitations have been well described (*32*, *33*). Taken together, while diet is assumed to be a major determinant of microbiome composition, an understanding of precise diet-microbiome interactions in humans under real-world disease and treatment conditions is lacking.

Patients with blood cancers who undergo allogeneic hematopoietic cell transplantation (allo-HCT) are typically hospitalized for several weeks while they receive chemotherapy, sometimes irradiation, and antibiotics. During this time, they exhibit drastic changes in nutritional intake (*34–36*) as well as severe microbiome injury (*4*, *37*) characterized by a loss of α-diversity and expansion of facultative anaerobe organisms. These microbiome shifts are associated with adverse clinical outcomes including bloodstream infections, graft-vs-host disease, and mortality (*37–40*). Disruptions to microbial ecology in this setting are driven by antibiotics (*4*, *20*, *41*, *42*) as well as intestinal inflammation induced by chemotherapy, irradiation and graft-versus-host disease (GVHD) (*39*, *43*). Here, we hypothesize that nutrition also contributes to microbiome dynamics. The weeks during which allo-HCT patients are hospitalized facilitates frequent collection of both stool specimens and high-precision dietary intake data; considering these data in light of these perturbations enables causal-inference approaches (*2*, *44*).

Thus, we hypothesized that the effect of diet can be quantified in the perturbed microbiome of allo-HCT patients to reveal mechanistic insights into diet-inducible microbiome modulation. For this, we collected daily real-time dietary intake data from allo-HCT recipients, paired with longitudinally collected fecal samples. We profiled fecal microbiome compositions and revealed associations between key microbiome features and dietary components. Intake of sweets during antibiotic exposure predicted low fecal microbiome α-diversity and expansion of *Enterococcus* abundance; and we validated this finding in a mouse model of post-antibiotic *Enterococcus* expansion.

## Results

Allo-HCT begins with an intensive course of chemotherapy that is intended to eliminate neoplastic cells and clear a niche for the transplanted allograft. These conditioning regimens can also cause painful mouth and throat lesions (mucositis), nausea, and poor appetite. Consequently, patients often have a poor diet and lose weight, during a period of increased caloric needs (*45*, *46*). These symptoms often persist until the graft is established in the bone marrow (engraftment) and produces new neutrophils—observed in this cohort after a median of 12 days (range 8-37). The severe degree of nutritional perturbation is underscored by the observation that a subset of patients in our cohort had such low dietary intake that they required nutrition support, either via intravenous infusion (total parenteral nutrition, TPN, n = 23 patients for a median of 11 days, range 2-63) or a nasogastric tube (enteral nutrition, EN, n = 5 patients for a median of 76 days, range 5-156).

### Longitudinal microbiota data paired with high-resolution nutritional intake data

We recorded the precise intake of food items consumed during 9,419 meals (**Fig. 1A**) consumed by 173 patients with blood cancers treated with allo-HCT between 2017-2022 (**Table 1**, range 8-128 meals/patient across −12-49 days). We mapped meal records to a hierarchical food nomenclature (Food and Nutrient Database for Dietary Studies, FNDDS) that classifies foods into nested categories (*47*), resulting in 9 distinct food groups (“Grain Products”, “Vegetables”, “Meat, Poultry, Fish, and mixtures”, “Sugars, Sweets, and Beverages”, “Milk and Milk Products”, “Fruits”, “Dry Beans, Peas, Other Legumes, Nuts, and Seeds”, “Fats, Oils, and Salad Dressings” and “Eggs”) in our data. A food taxonomy constructed from these categories (**Fig. 1B**) facilitated analysis of specific food types at various hierarchical aggregation levels as well as the application of metrics from ecology to summarize nutritional intake data (*11*). Thus, the dietary intake data in this study was take the form of, for example, “1 day prior to the fecal sample collected on day −5 relative to transplant day, the patient consumed 142 grams of baked, broiled or roasted chicken breast,” which is a distinct from the resolution offered by typical food frequency instruments (e.g., “chicken 2 times per week”) (*48*).

**Fig. 1.** Longitudinal microbiota data paired with high-resolution nutritional intake data. (**A**) Histogram of 9,419 meals recorded (top) and of 1,009 evaluable stool samples collected (bottom) from 173 patients during allo-HCT, where day 0 is the day of cell infusion. (**B**) Food tree of 623 unique food items according to the Food and Nutrient Database for Dietary Studies (FNDDS) nomenclature. The tree is colored by 9 broad food groups; tree levels are derived from numeric food-code hierarchies. The lengths of tick marks around the outer ring indicate the average per-meal consumption of each food item. (**C**) TaxUMAP visualization of recorded meals colored by the most consumed food group on that day. (**D-F**) Same TaxUMAP visualization colored by daily caloric intake (D), daily dietary α-diversity (E), and time relative to transplantation (F). (**G**) Daily caloric intake, where each point is a patient’s daily consumption. (**H**) Daily diet α-diversity (Faith’s phylogenetic distance). (**I**) Daily consumption of macronutrients: carbohydrates, sugars, fibers, protein, and fat. (**J**) Daily intake of the nine FNDDS food groups. (**K**) Fecal microbiome α-diversity (inverse Simpson index) of 1,009 stool samples. (G-K) lines: lines are LOESS averages (dietary variables in red, fecal microbiome variables in blue), shading indicates 95% confidence interval. (**L**) Scatterplot visualizing correlation between dietary intake and the microbiome on the same day (944 data points, 944 daily dietary intake data with the corresponding stool samples). The columns specify nutritional metrics including daily caloric intake and daily intake of the macronutrients carbohydrates, sugars, fibers, protein and fat. Plotted by row are natural-log transformed microbiome α-diversity and the log_10_-transformed relative abundances of the genera *Blautia* and *Enterococcus*. The blue lines denote linear regression lines; gray shading the 95% confidence level. The quantities ρ and p are Spearman correlation and corresponding p-value against the zero null, respectively.

To visualize the high-dimensional dietary dataset, we applied the TaxUMAP algorithm (*4*), a modification of UMAP that takes into account the taxonomic relationships between features, in this case food relationships defined by the FNDDS nomenclature (Fig. 1C-F). Each point represents a patient’s total food consumption on a single day. We color-coded dietary records by the most abundant food group consumed (based on imputed dehydrated mass) to reveal global patterns in consumption of meals dominated by certain high-level food groupings (**Fig. 1C**). Gradients across dietary TaxUMAP space were evident for total daily caloric intake (**Fig. 1D**), dietary α-diversity (**Fig. 1E**), and time relative to transplant day (**Fig. 1F**). This indicated that patients consumed more calories early during transplantation in the form of more complex meals, an observation confirmed in plots of time *vs.* daily caloric intake (**Fig. 1G**, p < 0.01) and dietary α-diversity (**Fig. 1H**, p < 0.01). Intake of the macronutrients carbohydrates, fat, fibers, protein and sugars all declined from hospitalization until day 12 (the median time to neutrophil engraftment in this cohort) (**Fig. 1I**, FDR q <0.001). The intake of all specific food-group items except for “sugars, sweets, and beverages” also declined over this time period (**Fig. 1J**, FDR q <0.01). Despite these global dietary trends, intra-and inter-patient variability in dietary intake were highly variable (**Fig. S2**), inspiring a detailed investigation of dietary association with microbiome injury patterns.

After filtering, we included 1,009 stool samples from a subset of 158 patients (median 5 samples per patient, **Fig. 1A, S1**) and profiled them by 16S rRNA gene sequencing. The decline in caloric intake and dietary complexity during allo-HCT corresponded to a decline in fecal microbiome α-diversity (**Fig. 1K**, p < 0.01), similar to that observed in several cohorts (*37*). Indeed, we found a significant correlation between fecal bacterial α-diversity and calorie consumption (**Fig. 1L, top left panels** p <0.01) as well as the magnitude of consumption of the macronutrients carbohydrates, sugar, fiber, protein, and fat (**Fig. 1L top row,** p <0.01 for each except sugar p = 0.04) in the 153 patients that had evaluable stool samples and nutritional intake data that were collected on the same day (**total n=944 days, median 5 days/patient**). Correspondingly, we observed positive associations between caloric (and macronutrient) intake and the relative abundance of the commensal genus *Blautia* (**Fig. 1L, middle row,** p <0.01 for each) which we previously associated with longer survival after allo-HCT and with lower rates of death from graft-*vs*-host disease (*39*). This genus is a member of family *Lachnospiraceae* that is a hallmark of a healthy human microbiome (*49*). Conversely, we observed inverse associations between caloric (and several macronutrients) intake and the abundance of *Enterococcus* (**Fig. 1L, bottom row,** p <0.01 for each, except sugar p = 0.06), a genus that includes several pathobionts that frequently cause antibiotic-resistant bloodstream infections and is associated with adverse outcomes following HCT, including graft-*vs*-host disease and mortality (*4*, *40*, *50*, *51*).

Clinical variables that might confound these correlations include the intensity of conditioning chemotherapy regimen (*43*) (**Table 1**) and antibiotic exposures. All 173 patients received at least one antibacterial antibiotic during their dietary collection period. They were typically treated initially with prophylactic antibiotics (fluoroquinolones and intravenous vancomycin (*52*)). Importantly, 138 of 173 (80%) patients also received broader-spectrum antibiotics during this time period, when they developed fever or other signs of potential infection (most commonly piperacillin-tazobactam, cefepime, linezolid, or a carbapenem for neutropenic fever or bloodstream infection; and metronidazole or oral vancomycin for *Clostridioides difficile* diarrhea). As these antibiotics are major drivers of dysbiosis in HCT (*4*, *20*, *41*, *42*, *53*) we took these exposures into account in our analysis.

### Consumption of sweets following antibiotic exposure predicts reduced fecal microbiota α-diversity

To quantify the contribution of dietary intake to microbiome composition in the context of such confounding clinical variables, we developed a Bayesian model that analyzes the relationship between microbiome composition and the dietary intake in the days preceding the collection of each fecal sample. We chose a dietary exposure period of two days preceding each fecal sample since variation in microbiome composition was best explained by windows of this duration in two different Procrustes analyses: one in which diet was summarized by macronutrient composition of meals (**Fig. 2A**, dashed line), or by named food-group items (**Fig. 2A**, solid line). Notably, two days was reported as the optimal width of a dietary window to correspond to microbiome composition among healthy volunteers (*11*).

**Fig. 2.** Consumption of sweets following antibiotic exposure predicts reduced fecal microbiota α-diversity. (**A**) Procrustes scores signify degree of correlation between average microbiome composition and dietary intake; dietary intake windows of 1, 2, 3, 4 or 5 days prior to each fecal sample were analyzed, either by macronutrient composition (dashed line) or by named food groups (solid line). The higher the Procrustes score, the better the correlation (**B**) Diagram representing the statistical model microbiome diversity or taxon abundances, which included antibiotic exposures; nutrition support modalities including total <u>p</u>arenteral (intravenous) <u>n</u>utrition, (TPN), and tube feeds referred to here as <u>e</u>nteral <u>n</u>utrition (EN) and dietary intake during the two days preceding the collection of each fecal sample (dashed line represents the interaction between antibiotics and food intake), as well as patient-level exposure (chemotherapeutic conditioning regimen intensity), and varying-effects terms (per patient and number of weeks spent in hospital). Blue boxes indicate time-varying quantities; the gray box indicates constants. (**C**) Posterior coefficient distributions of patient -level constants, namely the association of each conditioning intensity level with microbiome α-diversity. (**D**) Posterior distributions of association coefficients between predictors in the model and fecal microbiota α-diversity: the plotted coefficients estimate associations between consumption of 100 grams of each food group in the two days preceding fecal sample collection, as well the coefficients of interaction terms between each food group and antibiotic exposure, and additional fixed-effects predictor posteriors TPN, EN, and antibiotics independently of diet. (**E**) Posterior coefficient distributions from an analogous model using the prior two-day intake of 100 grams of macronutrients as dietary predictors; the protein category was omitted from the model because it was highly correlated with fat; (C-E) Dots, thick lines, and thin lines indicate the posterior medians, 66% CIs, and 95% CIs, respectively. 95% CIs not crossing zeros are highlighted in red. n=1,009 stool samples from 158 patients. (D-E) Rows on white backgrounds labeled in blue fonts are interaction terms between food and antibiotics; rows on light blue backgrounds labeled in black font are non-interactive terms. (**F**) Scatterplot visualizing the correlation between sweets consumption and natural log-transformed α-diversity stratified by antibiotic exposure during the preceding two days; lines from linear regression with 95% confidence level as shaded regions. The ρ and p values are Spearman correlations and corresponding p-value against the zero null, respectively. (**G**) Marginal effects plots of each food group consumption on the predicted ln(α-diversity) based on the different conditions: with or without antibiotics exposure in the prior two-day window while holding other variables constant. Lines represent the posterior predicted medians of marginal effects, shading represents the 95% interval. (F-G) Antibiotic-exposed and non-exposed groups are in red and blue, respectively.

The Bayesian model (summarized in **Fig. 2B**) includes treatment regimen (conditioning) intensity, nutrition-support events (TPN and EN), antibiotic exposure, and dietary intake during the prior two days as fixed-effect predictors of microbiome composition, as well as an interaction term between diet and antibiotic exposure, which captures an additional dietary effect on the microbiome when antibiotics are administered, profiling 1009 stool samples from 158 patients. Furthermore, as our data includes multiple timepoints per individual, the model includes varying-intercept terms for each patient (**Fig. S4**) and for the week relative to transplantation, capturing unmeasured differences between patients, as well as unmeasured exposures other than the explicit predictors.

With this model at hand, we then first quantified the associations of antibiotics, clinical parameters, and dietary food groups with microbiome α-diversity, measured by the inverse Simpson index, taking into account patient-specific effects. We found fecal samples from recipients of the mildest conditioning regimen (“Non-ablative”) had the highest average diversity (**Fig. 2C**), consistent with our prior report (*43*). As expected, exposure to antibiotics in the prior two days was inversely associated with bacterial diversity (**Fig. 2D**, “abx” term, median: −0.22, 95% credible interval, CI: (−0.39, −0.05) representing the highest posterior density interval). Although intake of “sugars, sweets, and beverages” (abbreviated here as “sweets”) alone was not obviously associated with fecal microbiome α-diversity, sweets intake in antibiotic-exposed samples was: each 100-gram increase in sweets intake was significantly associated with an additional median decrease of 0.28 natural-log units (95% CI: (−0.46, - 0.07)) of α-diversity, compared to samples without prior antibiotic exposure (**Fig. 2D**).

We also summarized dietary intake into macronutrient composition, which is independent of the FNDDS classification. Corroborating the association with sweets, intake of sugars during antibiotic exposure (“abx * Sugars” interaction term) was additive to the decline in diversity observed after sugar exposure alone (**Fig. 2E**) (median: −0.22, 95% CI: (−0.44, 0)); moreover, intake of the macronutrient sugars alone was not associated with worse microbiome injury (**Fig. 2E**). These associations were surprising because oral nutritional supplements, classified by FNDDS under the “sweets” food group (**Fig. S5**), are commonly recommended to transplant recipients (*54*, *55*). A similar result was observed in a variant of the model in which we considered analgesic use as an indicator of the most severely ill states of allo-HCT patients (**Fig. S6**); 52 (5.2%) fecal samples from 26 (15.0%) patients were exposed to the most intensive analgesics.

Having observed a relationship between microbiome injury and co-incident exposure to antibiotics and sweets, we asked if this was also apparent in the raw data, even without the adjustments for the many clinical variables encoded in the Bayesian model (**Fig. 2B**). Indeed, a strong inverse relationship was apparent between grams of sweets consumed and fecal microbiota α-diversity in samples from patients who were exposed to antibiotics (Spearman correlation −0.19, p < 0.01), but this relationship disappeared in the absence of antibiotic exposure (Spearman correlation 0.05, p −0.21) (**Fig. 2F**). To further explore the relationship between food groups, antibiotics, and microbiota injury in the multivariable Bayesian model, we inspected the marginal-effects plots (**Figs. 2G**). These revealed a prominent independent relationship between food intake and microbiota α-diversity only for sweets and only during periods of antibiotic exposure.

### Sugar and antibiotics synergize to exacerbate *Enterococcus* expansion in the gut microbiome of humans and in mice

We next asked whether sugar intake during antibiotic-induced microbiome perturbations might favor blooms of certain bacterial taxa (*37*). *Enterococcus* relative abundance had the most extreme negative correlation with α-diversity in this dataset (Spearman correlation rho: −0.32, FDR *P* < 10^−10^) (**Fig. 3A** and **Fig. S7**), consistent with our prior report that *Enterococcus* expansions are the most frequent injury pattern in this population (*51*). Here, using *Enterococcus* relative abundances (centered-log-ratio transformed) as the outcome variable in a modified version of the Bayesian model, sweets consumption during antibiotic exposure was associated with an additional increase of *Enterococcus* relative abundance (median: 0.67, 95% CI: (−0.01, 1.33)), compared to samples without prior antibiotic exposure (**Fig. 3B** and **Fig. S8**).

**Fig. 3.** Sugar and antibiotics synergize to exacerbate *Enterococcus* expansion in the gut microbiome of humans and in mice. (**A**) Spearman correlations between fecal microbiome α-diversity (Simpson’s reciprocal index) and the 5 significantly (adjusted p < 0.05) correlated genera with strongest correlations in either direction. The genera were pre-screened for presence in >10% of samples with >0.01% relative abundance. Blue and yellow represent negative and positive correlations between genera relative abundance and α-diversity, respectively. Other genera are plotted in **Fig. S7**. (**B**) Associations between *Enterococcus* relative abundances (centered-log-ratio (CLR) transformed) and antibiotic exposure (abx), total <u>p</u>arenteral (intravenous) <u>n</u>utrition (TPN), <u>e</u>nteral <u>n</u>utrition (EN), and intake of food groups in a variant of the Baysean model from Fig 2. Food group estimates are based on 100-gram intake in the two days preceding fecal sample collection. Box and whiskers indicate the posterior coefficient distributions of interactive associations. Dots, thick lines, and thin lines indicate the posterior medians, 66% CIs, and 95% CIs, respectively. n=1009 stool samples from 158 patients. Rows on white backgrounds labeled in blue fonts are interaction terms between food and antibiotics; rows on light blue backgrounds labeled in black font are non-interactive terms. For the interaction between sweets and antibiotics, 97.3% of the coefficient estimates were positive. A heatmap of results for other genera are shown in **Fig. S8**. (**C**) Sucrose exacerbates *Enterococcus* expansion following antibiotic treatment. *Upper panel*: fecal Enterococcal burden as measured by dilution plating on selective media; inset illustrates experimental setup. *Lower panel*: Enterococcal abundance from the top panel analyzed over time by trapezoidal area under the curve (AUC) to account for repeated measures from the same animals. 21-51 mice per treatment group; 2 independent experiments. The p-values were obtained by two-sided Wilcoxon ranked-sum tests (ns: not significant, *: P ≤ 0.05, **: P ≤ 0.01, ****: P ≤ 0.0001).

These observations in patients undergoing intensive cancer treatment suggested a model whereby antibiotic-induced microbiome injury, as manifest by α-diversity loss and pathobiont expansion, could be exacerbated by the consumption of sweets. To test this hypothesis, we established an experimental system in which microbiome disruption was induced in C57BL/6 mice with a single subcutaneous injection of the carbapenem antibiotic biapenem. This reproducibly induced a moderate expansion of endogenous Enterococci by 2-3 orders of magnitude that peaked at day 3 and largely resolved by day 6, as assessed by dilution plating on Enterococcal-selective agar (**Fig. 3C**, upper panel). As predicted by the analysis of patient data, supplementation of the standard chow diet with sucrose exacerbated the expansion of Enterococci on day 3 (median 3.53-fold, p = 0.049) and markedly sustained the Enterococcal expansion by a median of 7.30-fold by day 6 (p = 0.002). This effect of sucrose was also significant when accounting for repeated measures per mouse over time (p = 0.028, **Fig. 3C**, lower panel). Notably, sucrose treatment alone had no effect on Enterococcal burden in the absence of antibiotics. Taken together, our results provide observational evidence that consumption of dietary sugars, in the form of sweets food-group items, during or after antibiotic exposure decreases microbiome diversity and drives *Enterococcus* expansion, a novel microbiome-diet dynamic that we validated *in vivo*.

## Discussion

In this study, we analyzed a cohort of heavily treated patients whose fecal microbiome profiles and dietary intake were intensively monitored longitudinally. We found in patients with cancer undergoing allo-HCT that intake of foods enriched in simple carbohydrates (sugars) is linked to microbiome injury when the patients were concurrently exposed to antibiotics. These microbiome injuries—as observed in fecal samples with either low α-diversity or with relative expansion of *Enterococcus*—are clinically relevant as they have previously been reported to predict adverse clinical outcomes, including mortality, in several cohorts of allo-HCT recipients (*37*, *38*, *53*, *56*, *57*).

The dramatic perturbation experienced by the participants was key, as it facilitated analysis of how the microbiota responds to a profound stimulus. This “natural experiment” of longitudinal dynamics in a real-world clinically relevant setting offers opportunities for inference (*30*) that are distinct from those made in prior pioneering studies in which volunteers were profiled at presumed steady states (*10*, *11*), or assigned to specific diets (*13*, *15*, *17*, *58–60*). Additionally, the longitudinal design in which both fecal sampling and diet-intake were collected serially may increase statistical power for causal-inference hypotheses compared with cross-sectional sampling designs (*2*).

Collection of accurate dietary-intake data in humans is challenging, owing in part to the low resolution of food-frequency questionnaires and the human fallibilities in responding to recall survey instruments (*32*, *61*, *62*). The data-collection approach utilized here is not as rigorous as weighed food records; participants circled whether they ate 0, 25%, 50%, 75% or 100% of each item. This limitation, however, was mitigated by real-time meal logging and by the fact that the vast majority of meals were prepared by a large-volume hospital cafeteria with standardized portions and recipes, the underlying data of which were used in curating the dataset. Moreover, a dietician or their assistant also met with the patients thrice weekly to review their food diaries, motivate sustained participation with data collection, and clarify incomplete or inconsistent records, although this may have itself introduced bias through observer effects.(*63*)

Commonly used approaches to reduce the dimensionality of nutritional data include conventional macronutrient metrics (e.g., high fiber intake) and pre-defined diet-quality scores (*64*, *65*). However, associations between microbiota composition and such indices have not been consistently observed (*27*); here we looked beyond macronutrients and food-frequency habits by analyzing nearly-daily nutritional intake data considering specific food items (*11*).

Many variables in a clinically heterogeneous observational cohort can confound associations. The Bayesian analysis applied here accounted for several potentially confounding variables, including patient-level clinical variables such as conditioning intensity and temporal sample-level exposures such as antibiotics and parenteral nutrition (*66–68*). The inference from patient data that sugars and antibiotics may synergize to disrupt microbiome composition was corroborated as a causal effect in a mouse model of antibiotic-induced expansion of endogenous Enterococci that was exacerbated by sucrose supplementation. This is consistent with other reports that simple carbohydrates can induce dysbiosis and exacerbate experimental colitis or metabolic syndrome (*69–72*). Sucrose could have a positive effect on Enterococci expansion, or a suppressive effect colonization resistance that is usually conferred by other members of the microbiota community (*73*, *74*). That most simple carbohydrates are absorbed in the small intestine and only a minority typically reach the colonic microbiota raises a possibility that the sucrose modulates the host, which, in turn, shapes microbiota composition (*71*, *75*).

As low α-diversity and *Enterococcus* expansion have both been linked with adverse outcomes following allo-HCT (*37*, *51*), this study suggests that nutrition may have direct effect on patient outcomes via microbiome modulation; prospective clinical trials are needed sto guide nutritional recommendations. Moreover, avoiding sugar-enriched foods while taking antibiotics may have relevance in settings outside of allo-HCT. The general recommendation to limit sweets consumption (*76*, *77*) might, with further study, be tailored to abbreviated periods of avoidance to mitigate microbiome disruption during antimicrobial therapy.

**Fig. S1**. Flow of patients and sample selection through the study. Thrice weekly, patients who had recently been admitted for allo-HCT were invited to participate in dietary-intake collection and to donate fecal specimens on a biospecimen-collection protocol. ASV, amplicon sequence variants.

**Fig. S2**. Line plots illustrate the observed variation in the daily caloric intake, daily diet a-diversity (Faith’s phylogenetic distance), and fecal microbiota a-diversity (inverse Simpson index) across hospital stays. Each panel is one patient’s time course. The enlarged plot at the top left corner explains the details for every panel. The red, black, and blue line stands for the value of daily caloric intake, diet α-diversity and fecal α-diversity, respectively. The first two have values on the same numeric scale, therefore they share the same left Y axis. The microbiome α-diversity value is indicated by the Y axis on the right. The X axis represents the day relative to transplant. All panels share the same Y axis on both sides as the first one, with the X axis representing different temporal ranges of available data per patient. Alphanumeric codes are anonymized patient identifiers.

**Fig. S3.** Prior and posterior predictive checks of the Bayesian model (**Fig. 2B**) with natural-log-transformed fecal microbiome α-diversity as the outcome. (A) Prior predictive checks; the implemented priors are mildly regularizing and constrain the model (teal line) towards the domain of the observed diversity data (red histogram), in contrast to non-regularizing flat priors (black dashed line), which assume any outcome values as equally plausible *a priori*, and are shown for illustrative purposes. (B) Posterior predictive checks; observed data in red; ten independently simulated datasets drawn from the posterior predictive distribution are plotted in teal.

**Fig. S4**. Per-patient intercepts quantify the variation of microbiome α-diversity across patients. Each horizontal line represents a patient’s individual diversity fluctuations, as modeled in the Bayesian analysis. The thin whiskers indicate the 95% CI; thick whiskers the 66% CI. The dot signifies the median value. Red dashed line highlights if the interval is crossing zero. The patients are sorted by the median posterior coefficients.

**Fig. S5**. Effective per-meal average consumption of the top ten foods in gram weights in the “sugars, sweets, and beverages” group (referred throughout this manuscript as simply “sweets”). “Effective” refers to that only meals with that food’s consumption were included in the calculation. Dark pink bars denote the effective per-meal average consumption in dehydrated weight of each food in grams; the light pink bars represent the sugar content.

**Fig. S6**. Posterior distributions of association coefficients and patient-level differences in a variant of the main model that considered exposure to patient-controlled analgesia (PCA) opioids. The Bayesian model (**Fig. 2B**) included several variables to account for health status of the participants. However, mucositis might confound this analysis, as damage to the gastrointestinal lining can lead to translocation of luminal contents such as lipopolysaccharide (LPS), leading, in turn to fevers and to antibiotics; separately mucositis and effects of the conditioning can cause pain and nausea, leading to anorexia and recommendations by clinicians to consume oral nutritional supplements. We therefore conducted an expanded model in which we considered exposure to patient-controlled analgesia (PCA) opioids by intravenous pump in the two days preceding fecal-sample collection as an indicator of for severe mucositis and/or the overall severity of acute illness during the transplantation process. There were 52 (5.2%) fecal samples from 26 (15.0%) patients exposed. (**A**) Posterior coefficients of associations between 100 grams of food intake on its own in each group and with the exposure to antibiotics during the prior two-day window and bacterial α-diversity, as well as the association between exposure to TPN, EN, PCA, antibiotics and α-diversity. (**B**) Posterior distribution of the three levels of conditioning intensity with α-diversity. Dots, thick lines, and thin lines indicate the posterior medians, 66% CIs, and 95% CIs, respectively. 95% CIs not crossing zeros are highlighted in red.

**Fig. S7**. Spearman correlation between each genus relative abundance and the α-diversity of the stool sample (by inverse Simpson index). Genera observed in >10% of the samples at an abundance >0.01% were included. The p values from spearman correlation analyses were adjusted for multiple hypothesis testing via the Benjamini Hochberg method. Genera that met an FDR < 0.05 are included here. Blue bars indicate inverse correlation with diversity, while yellow bars positive correlation with diversity.

**Fig. S8**. Heatmap visualizing posterior association coefficients between food group consumption, nutritional support as well as antibiotics and CLR-transformed microbiome genus relative abundances. Genera were observed in >10% of the samples at >0.01% relative abundance were included in the analysis. Effects with 75% credible intervals (CI) to the right-hand side of zero, covering zero, and to the left-hand side of zero were displayed in red, white, and blue boxes, respectively. Effects with 95% CI not covering zero, 97.5% CI not covering zero, and 99% CI not covering zero were annotated by *, **, and ***, correspondingly.

## Methods

### Patients

Recipients of allo-HCT at Memorial Sloan Kettering Cancer Center between 2017 and 2022 who were consented to an IRB-approved biospecimen collection protocol, in accordance with the Declaration of Helsinki were eligible. Thrice weekly, we invited patients who had been recently admitted for allo-HCT to participate in dietary-intake collection. Neutrophil engraftment was defined as the first of three days of neutrophil count ≥500k/μl. Five patients died without achieving engraftment and were excluded from the analysis of median time to engraftment.

### Nutrition data collection and annotation

The hospital kitchen commercial computer system (Computrition, Bedford, MA) was configured to provide a printout that accompanied each meal tray to the bedside upon which patients were asked to indicate, immediately after each meal, whether they consumed 0, 25%, 50%, 75% or 100% of each item they ordered for that meal. These instruments were collected by a dietician or other research team members thrice weekly, during which missing entries were completed through informal bedside interviews along with encouragement to sustain motivation with the project. Consumption data were entered into the kitchen software, which was linked to the recipe and mass of each item. Data were manually vetted by a research dietitian for quality control (e.g., grossly implausible kilocalorie values resulting from sporadic typographic errors in the hospital kitchen records).

An eight-digit food code was assigned to each unique food item according to the classification of the Food and Nutrient Database for Dietary Studies (FNDDS) (*47*). The numeric codes are structures such that subsequent numeral positions differentiate foods within larger groups. For example, in the food code 14109010 for Swiss cheese, the first digit “1” denotes milk and milk products, the second digit “4” denotes cheeses, and each subsequent digit conveys progressively high-resolution classifications of foods. Water fractions from the FNDDS dataset were used to compute the dehydrated weight of the consumed food. For enteral nutrition, the listed water percentage in each formulation was used to calculate non-water volumes. The dehydrated weight was computed by converting the volume to grams based on 1.05 g/mL. The analyzed dataset thus listed the grams of FNDDS food items consumed in each meal on each day of hospitalization.

### Food tree construction

A food tree was constructed with the 622 unique FNDDS items consumed by the patients in this cohort, as done before (*11*)(https://github.com/knights-lab/Food_Tree). The tree spans nine broad FNDDS food groups, namely, Grain Products (abbreviated here as “grains”), vegetables, “Meat, Poultry, Fish, and mixtures” (abbreviated here at “meats”), “Milk and Milk Products” (abbreviated here as “milk”), “Sugars, Sweets, and Beverages” (abbreviated here as “sweets”), fruits, “Dry Beans, Peas, Other Legumes, Nuts, and Seeds” (abbreviated here as “legumes”), “Fats, Oils, and Salad Dressings” (abbreviated here as “fats”), and Eggs. In some cases, food items not explicitly classified in FNDDS contained ingredients from multiple categories (e.g., milk-based mango smoothie). We addressed this by manually categorizing foods based on which FNDDS food description they fit best to. For example, milk-based mango smoothie, despite having mango in it, fits the description of “11553110 fruit smoothie, with whole fruit and dairy”, therefore it was classified in the “Milk and Milk Products” group.

### Dietary data analysis: TaxUMAP and diet α-diversity

The hierarchical organization of the FNDDS vocabulary facilitated application of α-diversities to diet data using Faith’s phylogenetic distance (*78*), which was implemented in Qiime2 (qiime2-2021.11)(*79*) with the “qiime diversity alpha-phylogenetic” function with the faith_pd metric. The food tree taxonomy was utilized, as well as the dehydrated weight consumption of the food item represented by food code per patient per day. The TaxUMAP method (*4*) was used to visualize compositional similarities between the patients’ daily meals, similar to beta diversity. The fraction of each consumed food represented by a food code per patient per day was used to calculate the food tree taxonomy.

### Fecal microbiome analysis

Fecal sample inclusion criteria and flow through the study are detailed in **Fig. S1**. Microbiome profiling by 16S rRNA sequencing was performed as described (*80*). Briefly, bacterial cell walls were disrupted and nucleic acids isolated using silica bead-beating and phenol-chloroform extraction, and the V4-V5 variable region of the 16S rRNA gene was amplified. Amplicons were purified either using a Qiagen PCR Purification Kit (Qiagen, USA) or AMPure magnetic beads (Beckman Coulter, USA) and quantified using a Tapestation instrument (Agilent, USA). DNA was pooled to equal final concentrations for each sample and then sequenced on the Illumina platform. The median read count was 132,978 and the range was from 1,196 to 8,698,524. The 16S sequencing data were analyzed using the R package DADA2 (version 1.16.0) pipeline with default parameters except for maxEE=2 and truncQ=2 in filterandtrim() function (*81*), 16S FASTQ files were capped at 10^5^ reads per sample. Amplicon sequence variants (ASVs) were annotated according to NCBI 16S database using BLAST (*82*). Microbiome α-diversity was evaluated using the inverse Simpson index, a summary statistic of both the richness and evenness of the bacterial community. Taxa abundances were summarized at the genus level. Trends in microbiome and nutrition dynamics over time (Fig. 1G-J) were analyzed by generalized estimating equations using the geeglm function in geepack (1.3.9) in R.

### Procrustes test

Since we collected serial fecal samples and serial dietary intake data, a question arose as to how many prior days of dietary intake to correlate with any given fecal sample. To find the optimal time window of dietary intake to consider, we assessed concordance between microbiome and dietary datasets by Procrustes analysis, as done previously (*11*). We considered dietary data in two alternative intake metrics, first by macronutrients and second by specific named foods in FNDDS categories. Principal Coordinate Analysis (PCoA) was conducted in Qiime2 (qiime2-2021.11) by first converting the counts data to biom format, then to qza format, then for stool samples the Bray-Curtis distance was used for the PCoA analysis with the macronutrient data. For food-code data, the unweighted unifrac distance was used to calculate beta diversity PCoA with the food taxonomy information. The resulting principle coordinates were incorporated to compute a sum of square value using the Procrustes function from the vegan (2.5-7) package (*83*). A Procrustes score was defined as the difference between the minimal sum-of-squares from the five tested scenarios and the corresponding one scenario was computed.

### Bayesian multilevel model

#### Data preparation

For macronutrient analysis, for each stool sample included the dietary data were summarized as the previous two-day average intake of sugars, fibers and fat in grams. For the food-group analysis, for each stool sample included the dietary data were summarized as the previous two-day average intake of grains, vegetables, meats, milk, sweets, fruits, legumes, fats and eggs. Gram weights were divided by 100 so that the resulting coefficients represent the expected change in the outcome variable per each 100-gram intake of the dietary component. Conditioning intensity was a three-level factor variable comprising (in increasing order of intensity) nonmyeloablative (“nonablative”), reduced intensity conditioning (“reduced”), and myeloablative (“ablative”). EN, TPN, and patient-controlled analgesia were binary variables encoded as true if the patient was exposed in the two days preceding fecal collection and otherwise false. Likewise, if the stool sample was collected after exposure to microbiome-damaging antibiotics in the two-day window before it, it was set to true; otherwise, false. The microbiome-perturbing antibiotics considered were piperacillin/tazobactam, carbapenems, cefepime, linezolid, oral vancomycin, and metronidazole; these were most commonly used as empiric therapy (most commonly for neutropenic fever) or in a pathogen-directed fashion (most commonly for bloodstream infections or *C. difficile* diarrhea) (*37*). Prophylactic antibiotics fluoroquinolones (*84*, *85*) and intravenous vancomycin (*52*, *86*) were not considered.

Association of foods and antibiotics with microbiome diversity were also analyzed as interaction terms, with the assumption that food correlations with microbiome may be different depending on whether the patients had antibiotics in the prior two-day window or not. We considered repeated measurements and thus per-patient differences in microbiome features across samples by implementing a per-patient varying intercept. We also considered additional exposures during hospitalization that we did not capture explicitly, which could therefore confound the analyses, and therefore also accounted for time by grouping samples taken during similar timepoints along the therapy course into weekly categorical time bins, as done before (*87*, *88*), which were included in the model as varying intercepts. Our reasoning for this implementation was that we lack a biologically informed prior model for the effect of time on the microbiome over the short course of several weeks, and because the *a priori* known causal effectors of microbiome changes such as antibiotic exposures and diet were explicitly included in the model. We partially pooled the data via a varying-effects implementation because we believe that while the microbiome could vary over time due to unmeasured confounders during the highly planned and controlled HCT treatment regimens, the time bins do contain mutual information justifying their crosstalk during Bayesian inference. The time bins were implemented in the format of weeks relative to transplant; for example, the sampling window expressed in days as [−7,0) is the week before transplant, and [7,14) the second week after transplant.

When the outcome was microbiome α-diversity, it was analyzed as the natural-log-transformed inverse Simpson’s index. When the outcome was genus abundance, it was analyzed as centered log ratio (CLR)-transformed raw ASV count of the genus after adding a pseudo-count of 0.5 reads with the clr function in the compositions package (2.0-6) (*89*). Ninety genera were included in the analysis since each genus was detected with a relative abundance greater than 0.01% in at least 10% of the samples.

#### Model construction, sampling and results visualization

brms(2.16.3)(*90–92*) and rstan(2.26.4)(*93*) packages were used to build and run the model, with the model formula:

~~~
log(simpson_reciprocal) ∼ 0 + ave_fiber + ave_fat + ave_Sugars + ave_fiber:abx +
ave_fat:abx + ave_Sugars:abx + intensity + EN + TPN + abx + (1 | mrn) + (1 | timebin)
~~~

for nutritional intake represented as the macronutrients, or:

~~~
log(simpson_reciprocal) ∼ 0 + ave_fruit + ave_meat+ ave_milk+ ave_oils+ ave_egg+
ave_grain+ ave_sweets+ ave_legume+ ave_veggie + ave_fruit:abx + ave_meat:abx +
ave_milk:abx + ave_oils:abx + ave_egg:abx + ave_grain:abx + ave_sweets:abx +
ave_legume:abx + ave_veggie:abx+ intensity + EN + TPN + abx + (1 | mrn) + (1 |
timebin)
~~~

for nutritional intake represented as the food groups.

We chose regularizing, mildly informative priors to avoid overfitting (**Fig. S3**); specifically, three levels of the conditioning intensity were used as intercept terms with normal distributions with mean 2 and standard deviation 0.1 as priors, intercept food group effects in units of per 100g with or without antibiotic exposure were set to normal distributions with mean zero and standard deviation 1; the coefficient for the binary indicator for antibiotic exposure was set to a normal distributions with mean zero and standard deviation 0.5; EN and TPN exposure were set to normal distributions with mean zero and standard deviation 0.1. Priors were chosen based on prior predictive simulations so as to constrain the model towards plausible outcome domains (**Fig. S3A**). The inference parameters were set to “warmup = 1000, iter = 3000, control = list(adapt_delta = 0.99), cores = 16, chains = 2, seed = 123”, which means the model will sample 1000 steps during warm up, and 3000 iterations in two chains, adapt_delta is raised to 0.99 instead of 0.8 to avoid divergent transitions. The model inference can be reproduced in full via the model code and the processed data deposited publicly via GitHub repository: https://github.com/mskcc-microbiome/Nutrition_microbiome.

To quantify the effect of dietary predictors in concert, we leveraged the full posterior to conduct marginal effects analyses (**Fig. 2G**) as well as posterior predictive checks (**Fig. S3B**). Prior and posterior predictions were visualized with ggplot (3.3.5)(*94*), tidybayes (3.0.2)(*95*) and ggpubr (0.4.0) packages. The 66% and 95% CIs are shown as thicker and thinner lines on the coefficient plots, while the posterior medians are shown as dots.

When modeling taxon abundance, the formula was changed to:

~~~
CLR(taxon) ∼ 0 + ave_fruit + ave_meat+ ave_milk+ ave_oils+ ave_egg+ ave_grain+
ave_sweets+ ave_legume+ ave_veggie + ave_fruit:abx + ave_meat:abx + ave_milk:abx +
ave_oils:abx + ave_egg:abx + ave_grain:abx + ave_sweets:abx + ave_legume:abx +
ave_veggie:abx+ intensity + EN + TPN + abx + (1 | mrn) + (1 | timebin)
~~~

with the same running parameters.

The posterior coefficient distribution for the 90 investigated genera were summarized in a heatmap (**Fig. S8**) using ggplot. The marginal effects of each food group consumption on the predicted diversity are calculated with the conditional_effects function from brms package. The method used in the function was “posterior_epred”. While visualizing the predicted diversity as the intake of one food group changes, the rest of the food groups intake are held at average level, with intensity of the conditioning regimen set to be “nonablative” with no TPN or EN exposure.

### Mice

Female C57BL/6 mice aged 6-8 weeks procured from rooms RB03 or RB04 at Jackson Laboratory. Mice were single-housed in shoebox-style cages for 2-3 days before each experiment.

### Antibiotic intervention and Diet Preparation

Mice were treated with a single subcutaneous injection of biapenem (2mg in 100μl phosphate-buffered saline) or vehicle. In addition to standard mouse chow, the animals were fed Hydrogel cups HydroGel cups (ClearH2O; Cat: 70-01-502) with or without 5% added sucrose (sucrose diet and control diet, respectively). Cups were replenished every 48 hours. The experimental groups were control diet + vehicle, control diet + antibiotic, sucrose diet + vehicle, and sucrose diet + antibiotic, consisting of a total of N = 21, 39, 21 and 51 mice per group, respectively, across two independent experiments.

### Stool collection and colony counting

Fecal pellets were collected in a biosafety hood directly from the mouse into a barcoded pre-weight sterile tube and kept on ice until homogenized in 1mL of PBS and serially diluted. Twenty microliters of each dilution were plated on *Enterococcus*-selective agar plates (BD Cat: 212205) and incubated for 48 hours at 37°C under ambient air. Colonies from a plate with an enumerable density of colonies were counted manually.

### Trapezoidal AUC

The trapezoidal AUC was calculated between day 0 and day 3, as well as day 3 and day 6, respectively, for each mouse in the experimental setting, following the trapezoidal rule. The day 0 raw count was subtracted from day 3 and day 6 for each mouse before applying the trapezoidal rule. The reported total trapezoidal AUCs for each treatment group were the sums of the above two separate time periods and they were compared with the Wilcoxon rank sum test.

## Supporting information

Data S1 NCBI accession sampleid

Data S2 patients nutrition intake

Data S3 nutrition microbiome correlation

F1 overview raw 072 new

F2 model results 176 new

F3 current 178 tsoni

S1 stool sample selection

S2 all patients timecourse 1 085

S2 all patients timecourse 2 085

S2 all patients timecourse 3 085

S2 all patients timecourse 4 085

S3 prior and post pred 173

S4 mrn intercepts forest 179

S5 top10 eaten foods of sweets 099

S6 pca diversity 084

S7 genus diversity correlation 178

S8 genus heatmap 180 tsoni

Table 1 patient summary 003

## Author contributions

K.A.M., M.L.B., R.R.J., M.R.M.B., J.S. and J.U.P. conceptualized the project; A.D., P.A.A., M.R., B.G., S.R., E.H., M.L.B., A.G., J.B.S., A.G.C., D.G.B. and P.A.G. collected data; A.D., T.F., W.P.J., T.F., N.R.W., C.D., C.Z., F.M., A.P.S., A.L.C.G., A.J.J., D.K., M.R.M.B., J.S. and J.U.P. were involved in data analysis; A.D., P.A.A., T.F., W.P.J., T.F., N.R.W., M.R., B.G., S.R., E.H., M.B., C.Z.B., M.L.B., T.P., A.G., L.A. A., C.D., C.Z., F.M., A.P.S., J.B.S., A.G.C., D.G.B., P.A.G., A.L.C.G., A.J.J., D.K., R.R.J., J.S. and J.U.P. contributed to methodology; M.R.M.B. and J.U.P. helped with funding acquisition; A.G., J.B.S., M.P., S.A.G., M.R.M.B. and J.U.P. helped with project administration; M.R.M.B., J.S. and J.U.P. supervised the project; A.D., J.S. and J.U.P. wrote the original draft; K.A.M., O.M., M.R.M.B., J.S. and J.U.P. helped with the review and editing of the draft.

## Data and materials availability

the code used in the analysis can be found at https://github.com/vdblab/nutrition_and_microbiome. The amplicon sequencing data have been uploaded to NCBI, and the accession number of the samples are included in Data S1.

Data S1. A table with sample IDs and the accession numbers of the amplicon sequencing files.

Data S2. Anonymized patients detailed nutritional intake data including the foods name, eaten portion total calories and individual macronutrients gram weight on each hospital day.

Data S3. A table with summarized information of the average intake of each food group during the prior two days of each stool sample collection, which was used in the Bayesian model.

