## Supplementary figures and images for "Sugar-rich foods exacerbate antibiotic-induced microbiome injury"

### F1 overview raw 072 new

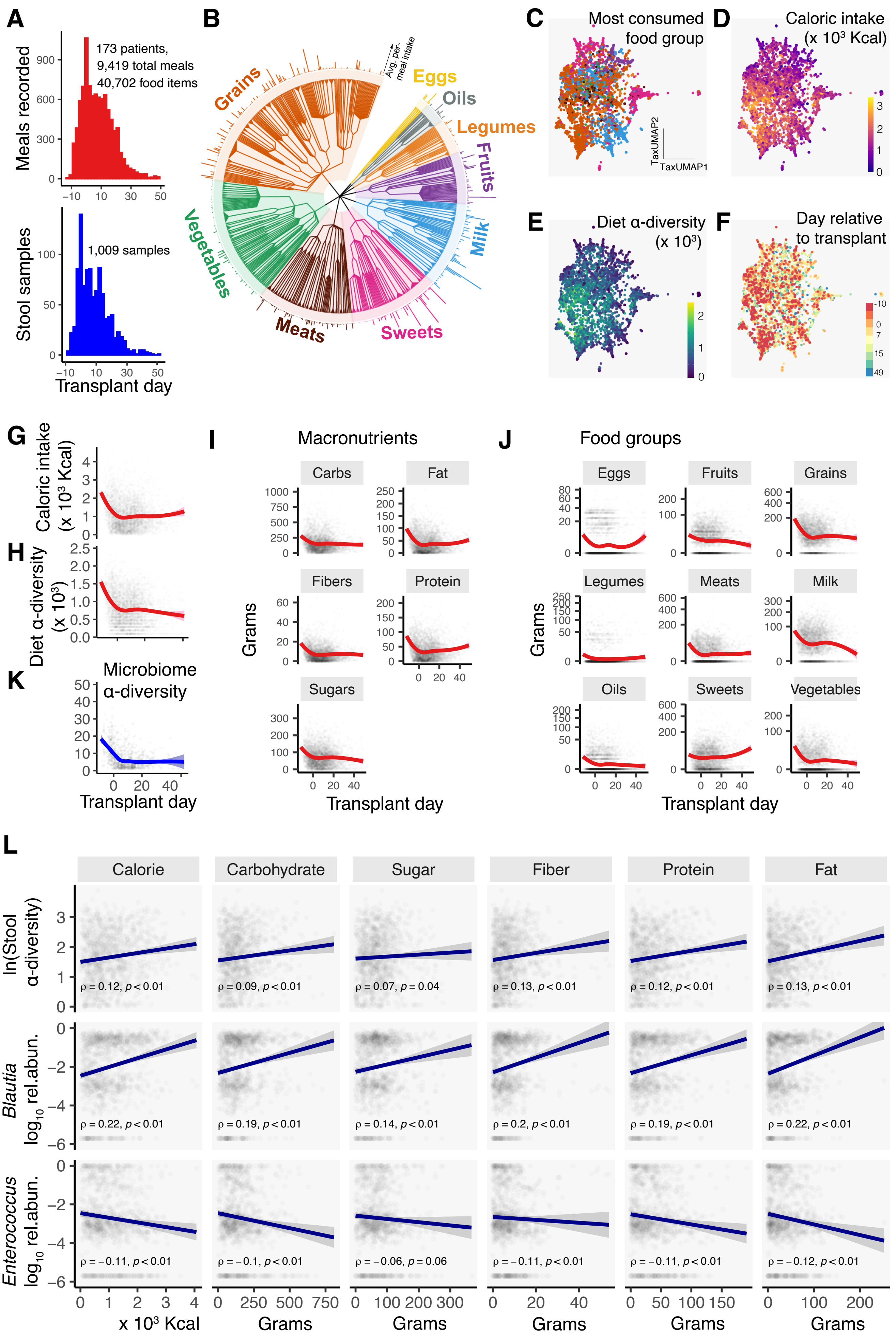

### F2 model results 176 new

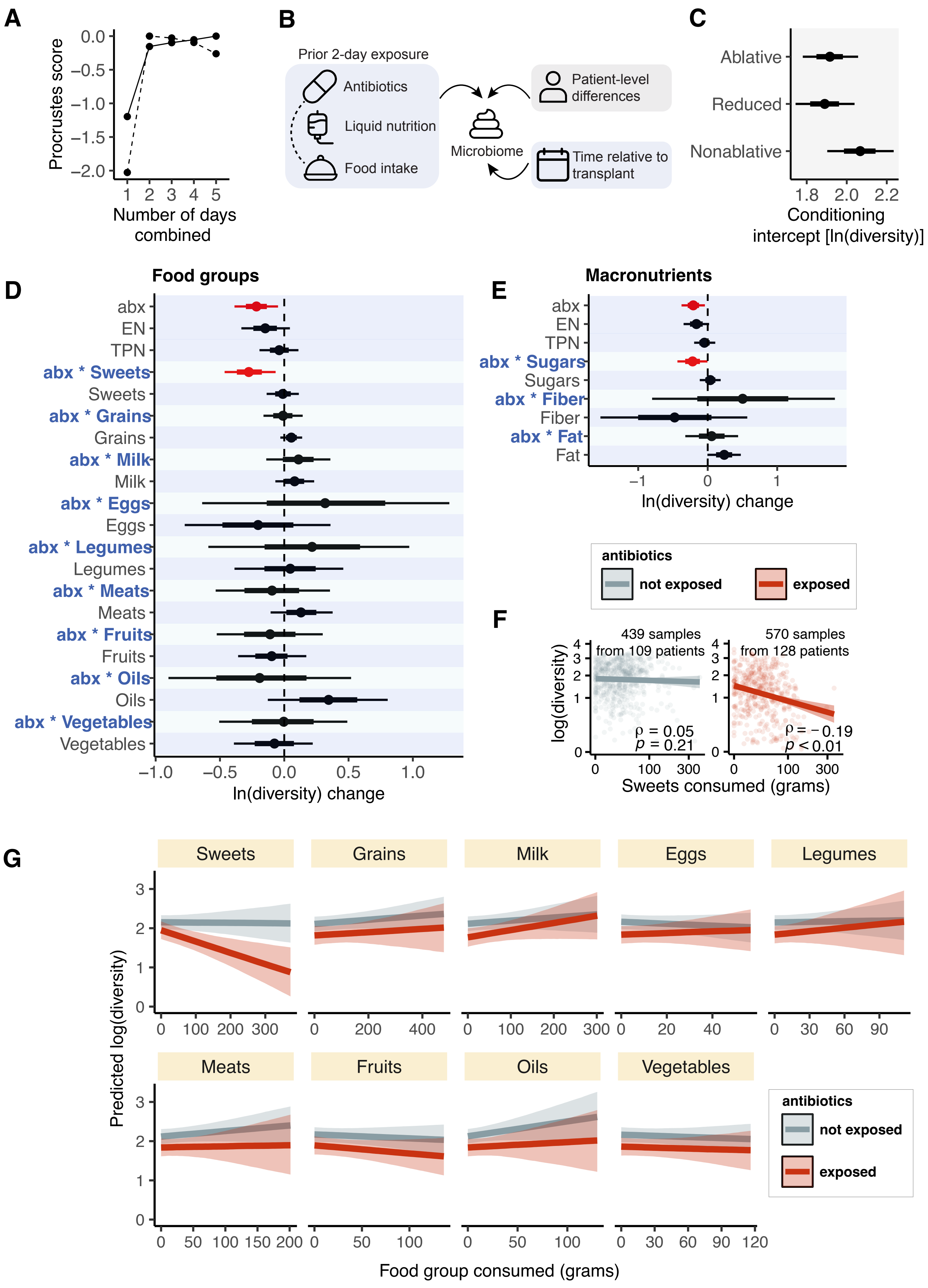

### F3 current 178 tsoni

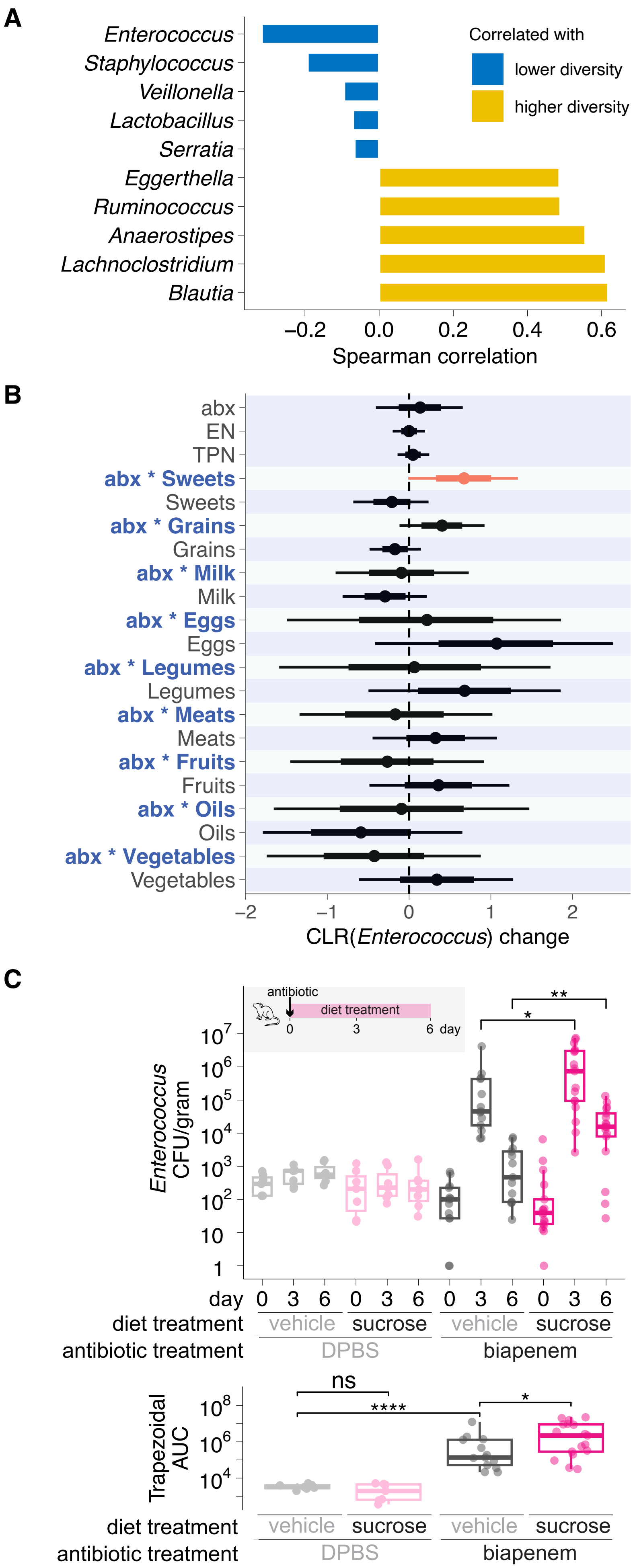

### S1 stool sample selection

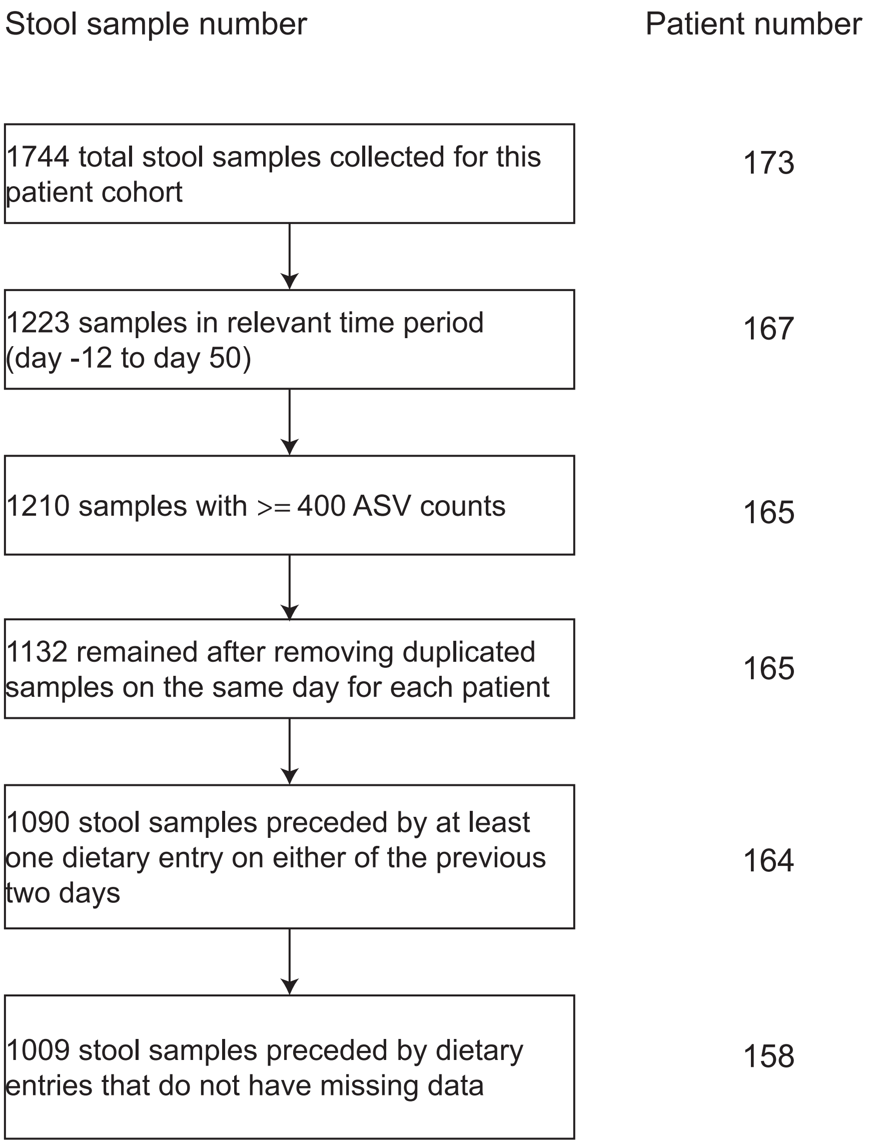

### S2 all patients timecourse 1 085

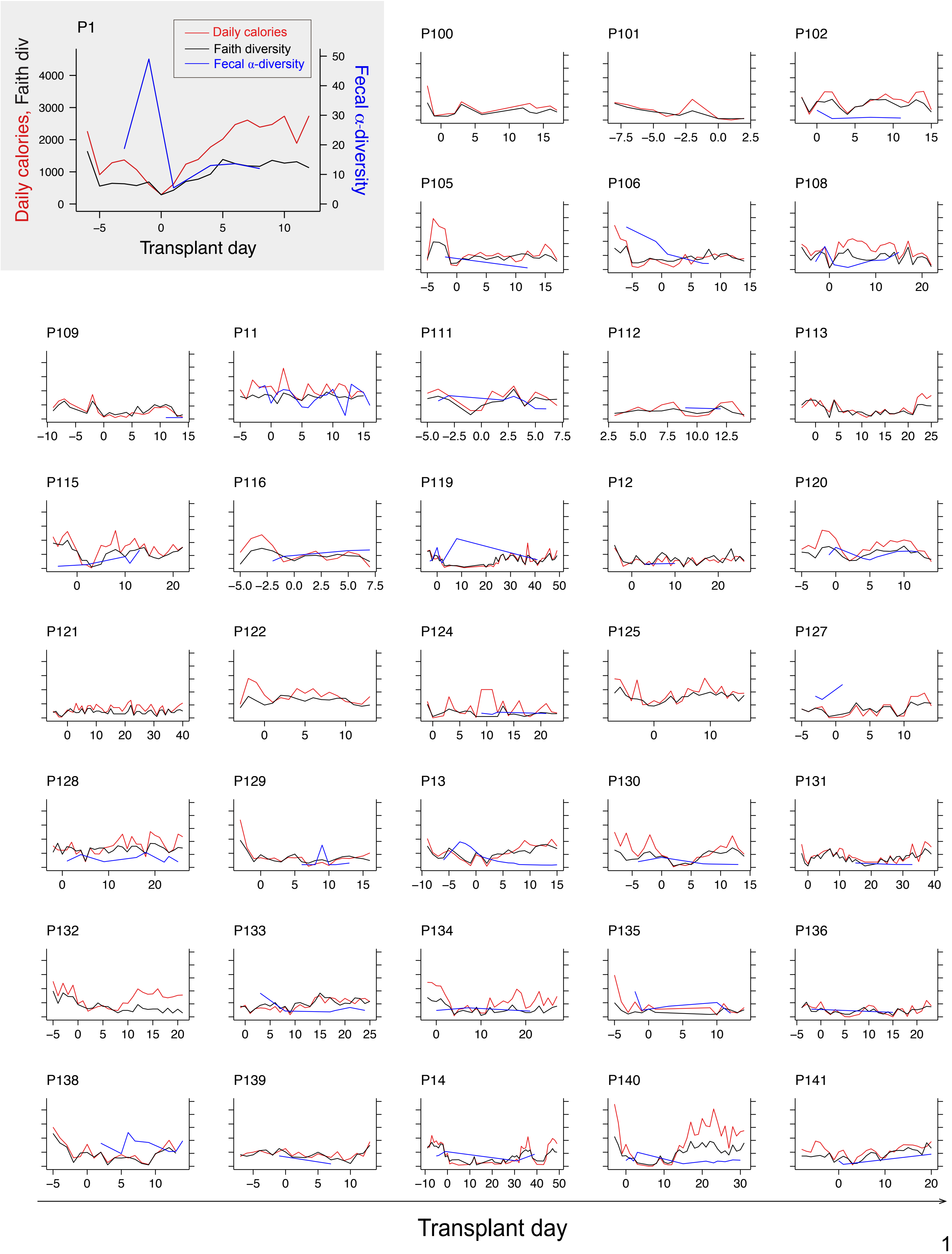

### S2 all patients timecourse 2 085

#### Fig. S2 (Continued)

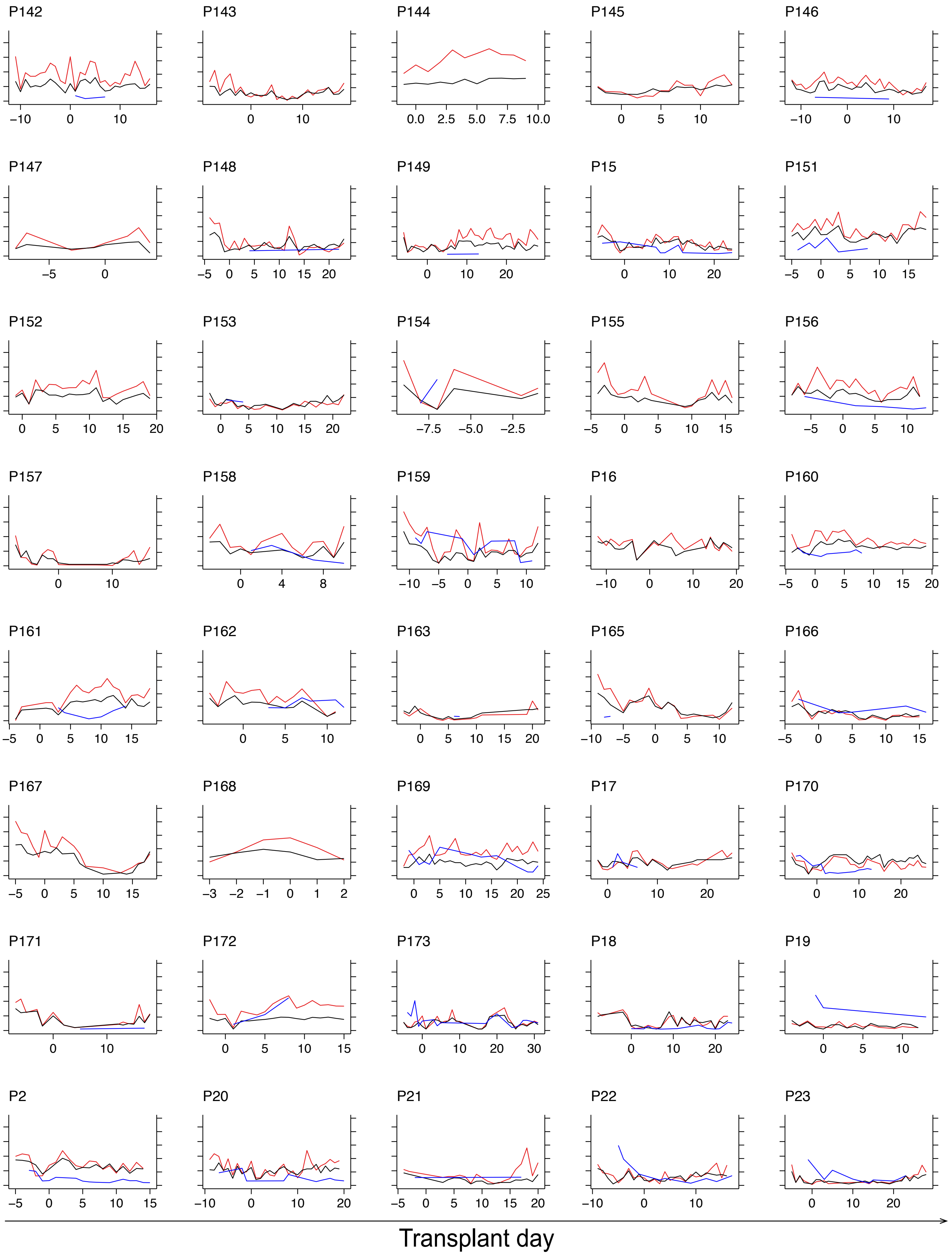

### S2 all patients timecourse 3 085

#### Fig. S2 (Continued)

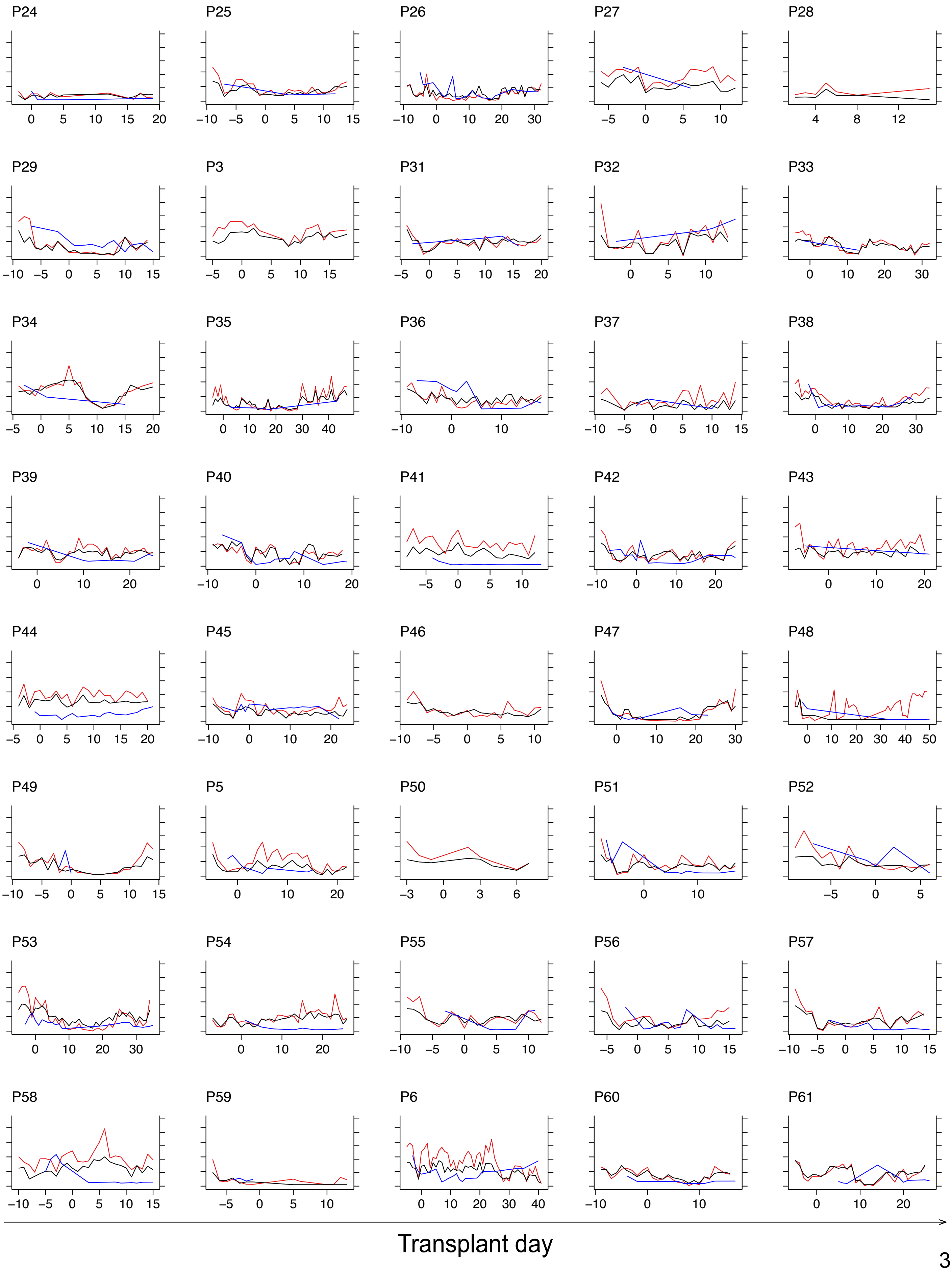

### S2 all patients timecourse 4 085

##### Fig. S2 (Continued)

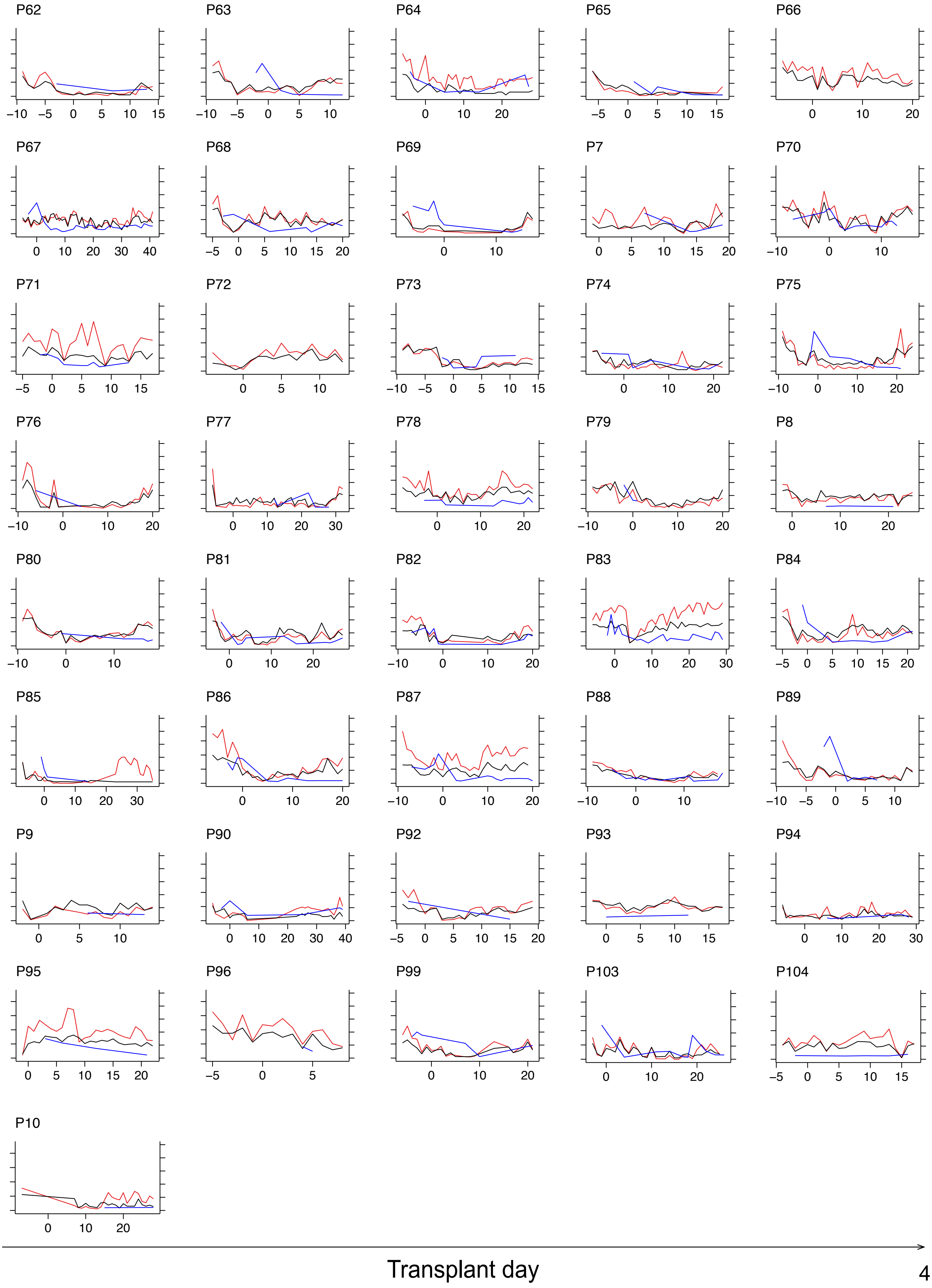

### S3 prior and post pred 173

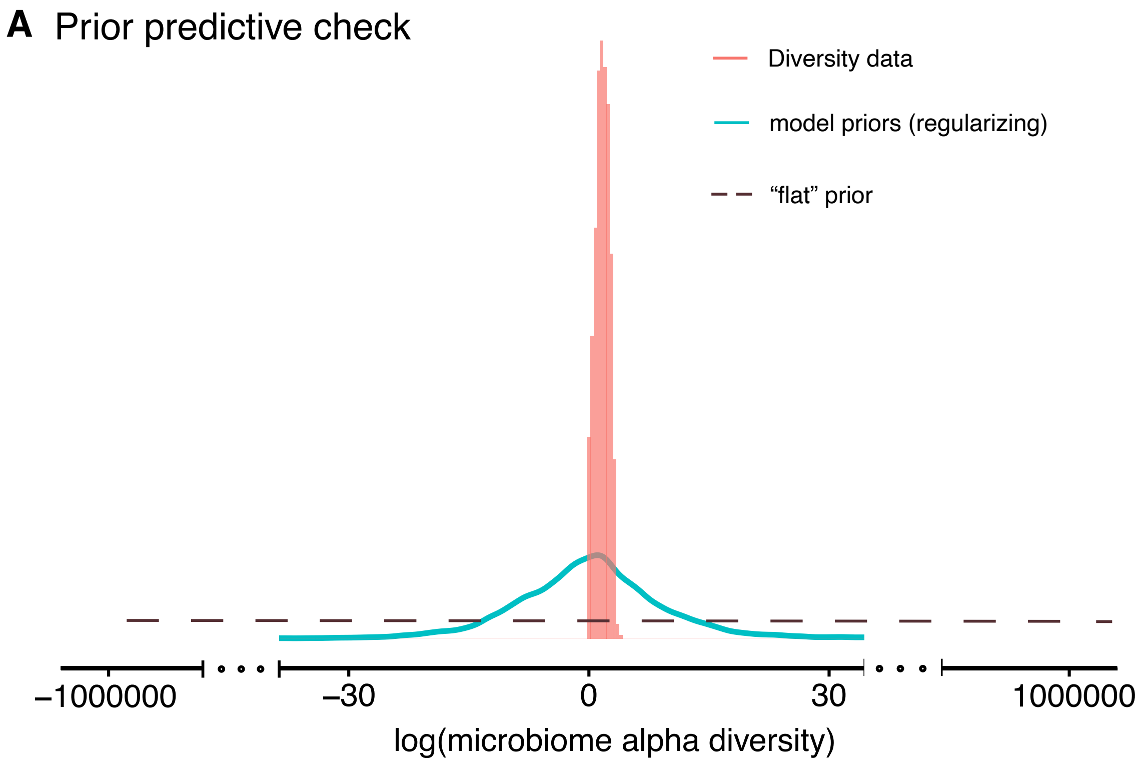

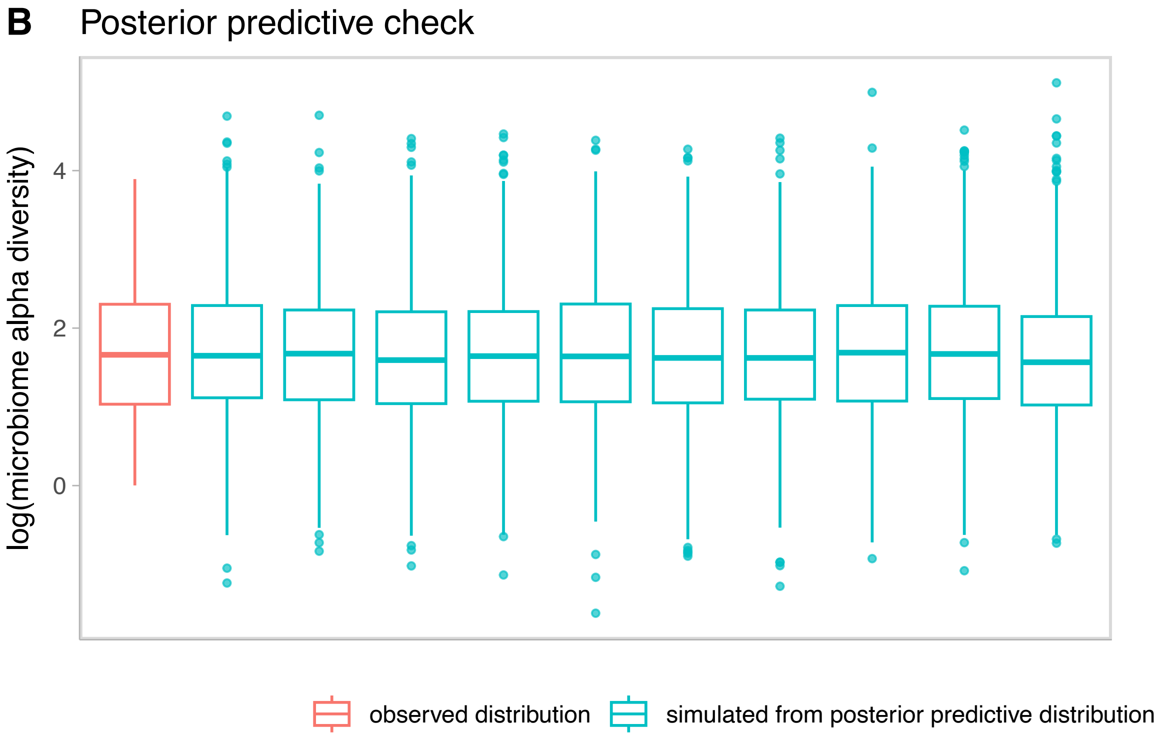

### S4 mrn intercepts forest 179

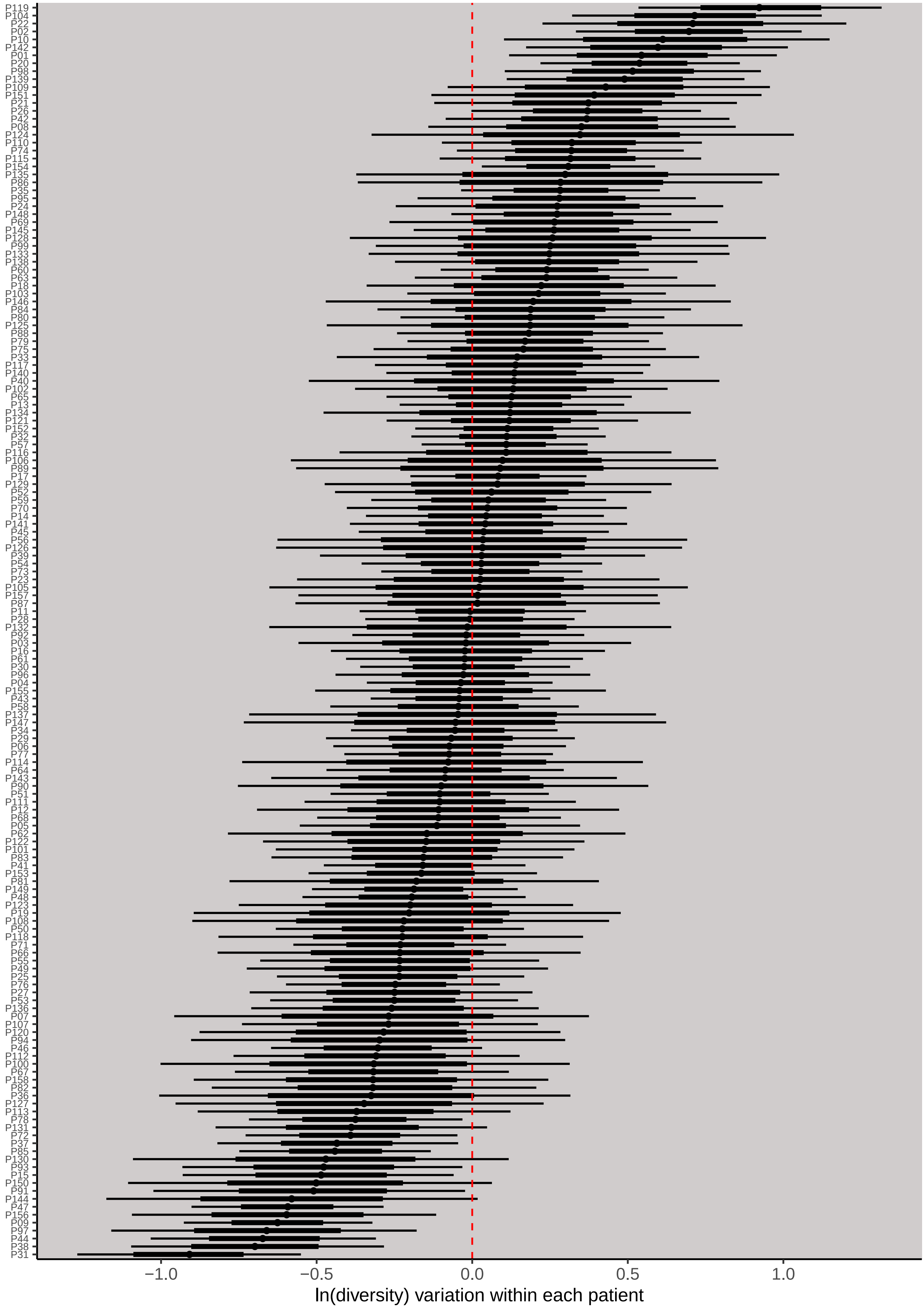

### S5 top10 eaten foods of sweets 099

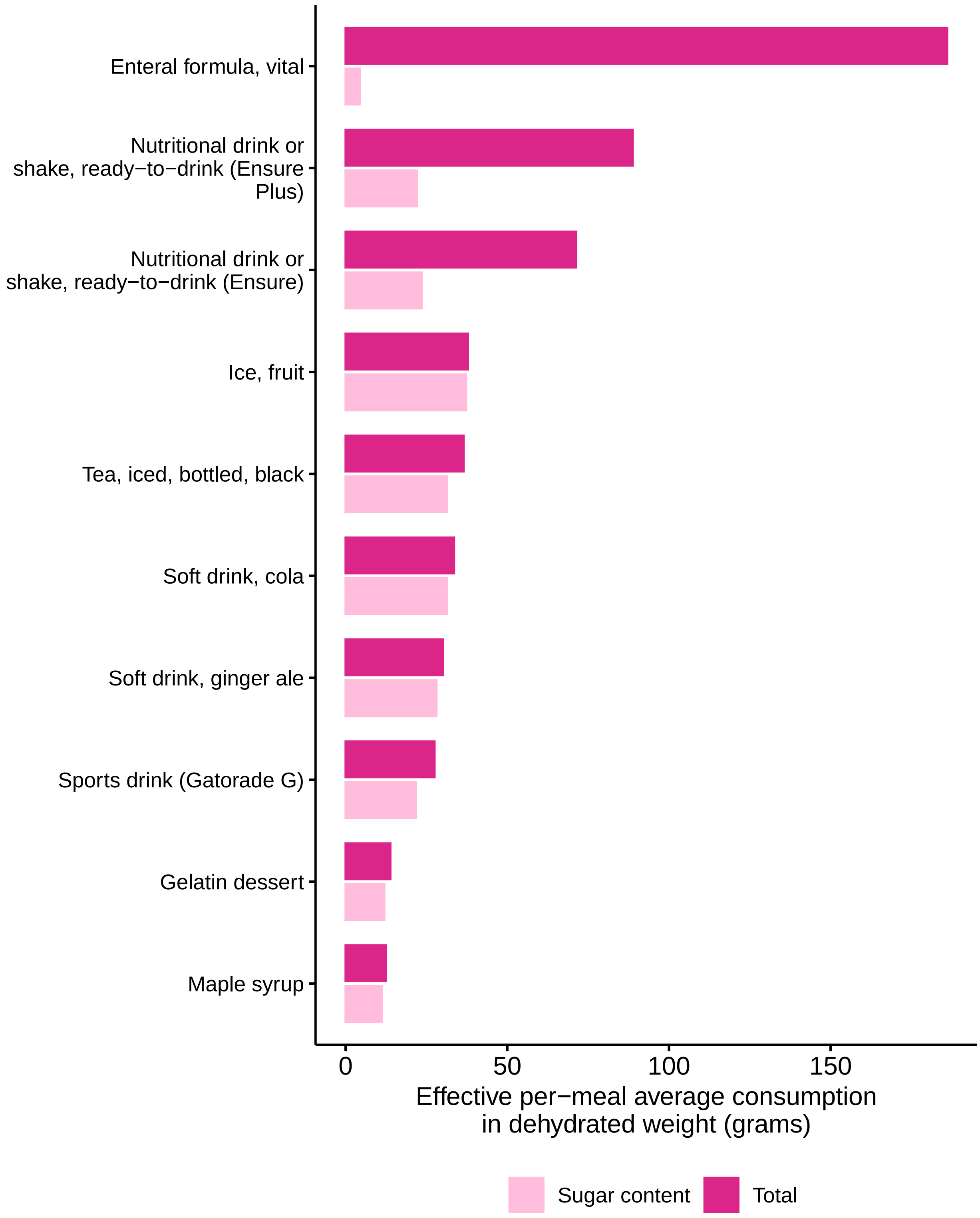

### S6 pca diversity 084

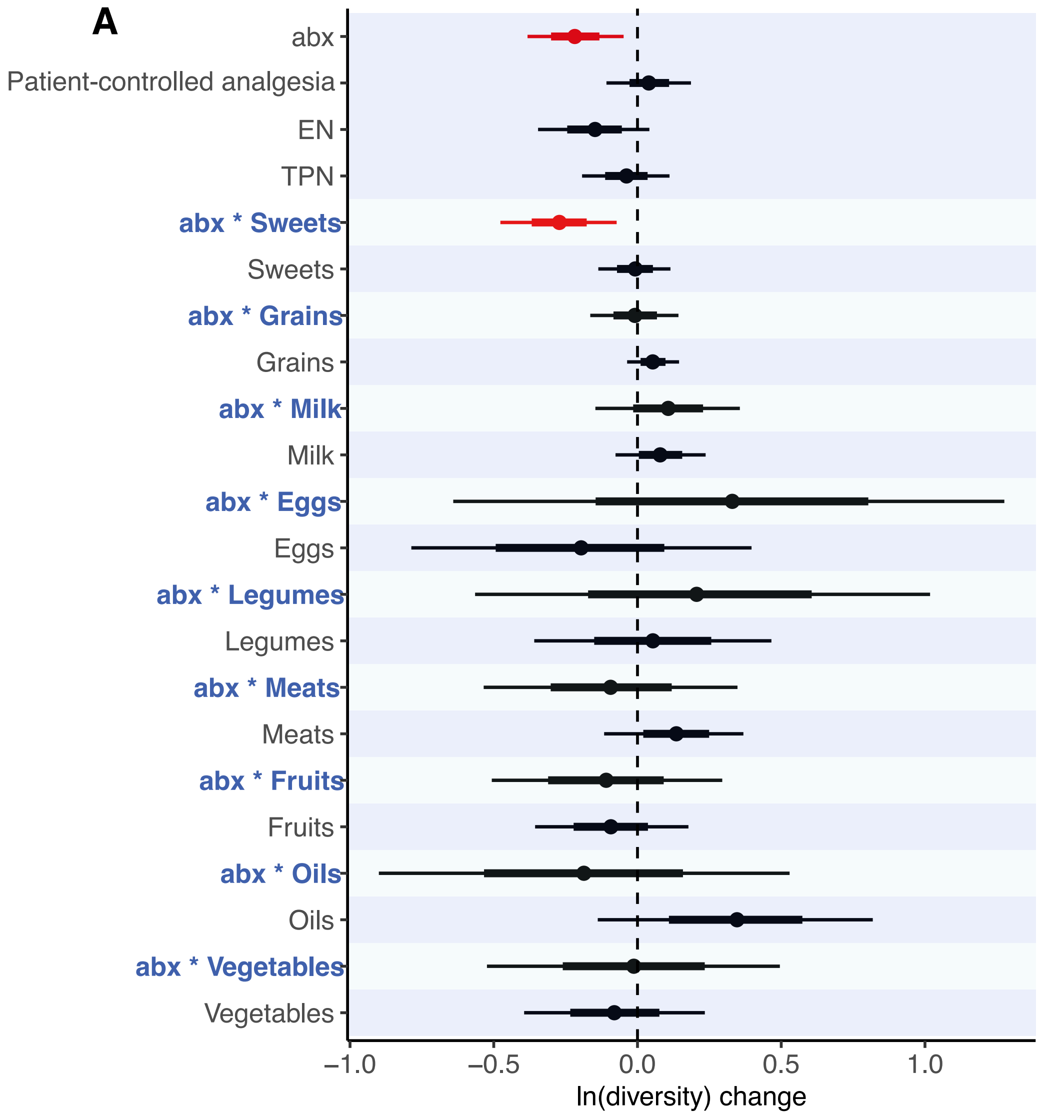

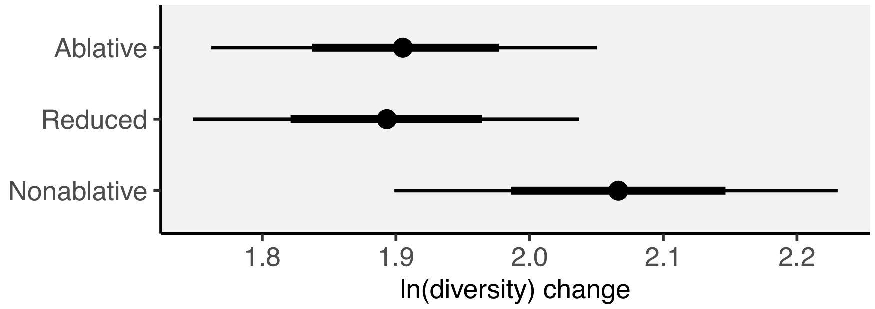

### S7 genus diversity correlation 178

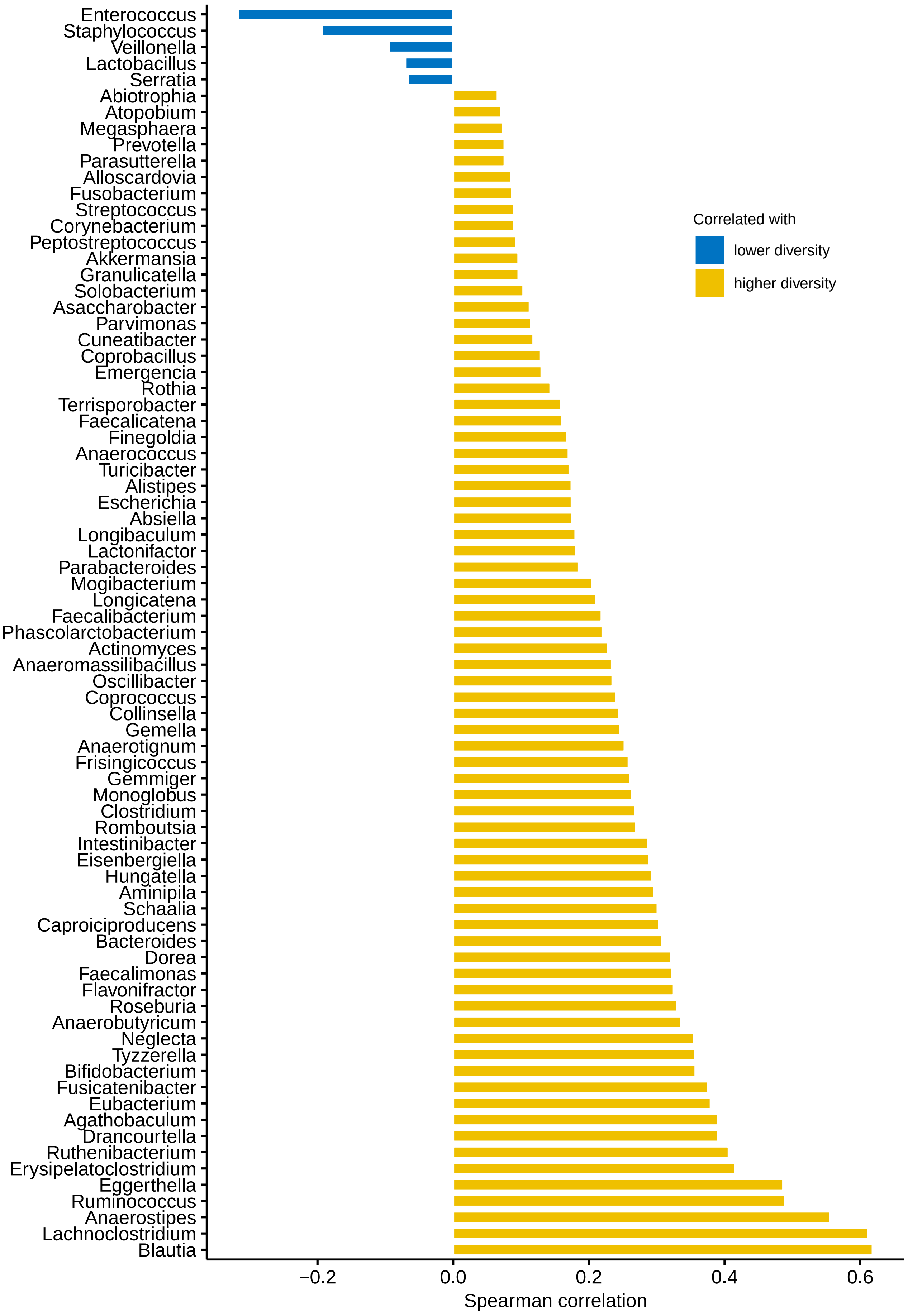

### S8 genus heatmap 180 tsoni

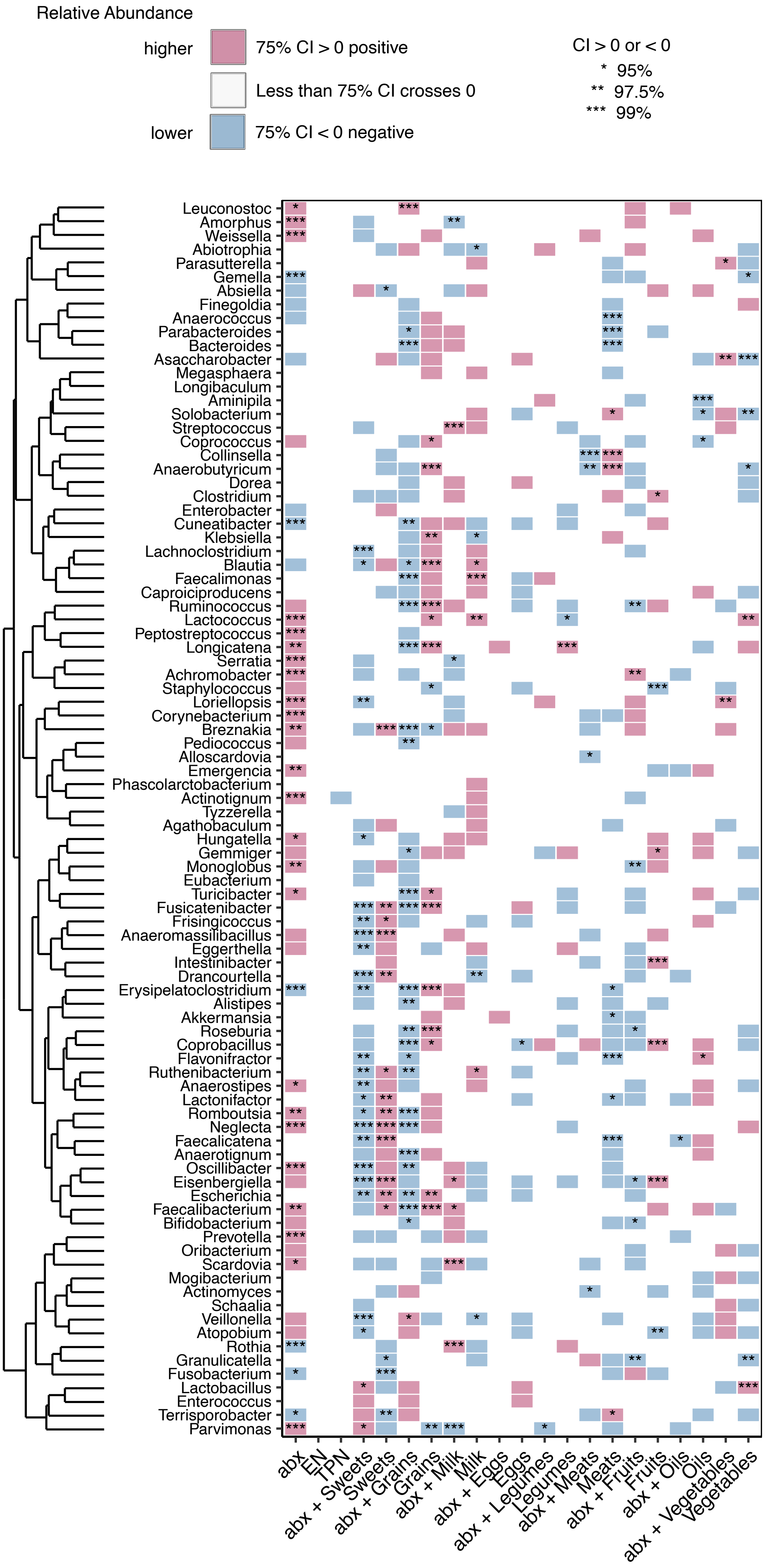
