## Supplementary material for "Sugar-rich foods exacerbate antibiotic-induced microbiome injury": Table 1 patient summary 003

| Characteristic | N = 173[1] |
|---|---|
| **Age** | 58 (12) |
| **Sex** | |
| Male | 95 (55%) |
| Female | 78 (45%) |
| **Disease** | |
| Acute myeloid leukemia | 68 (39%) |
| MDS/MPN [2] | 57 (33%) |
| Non-Hodgkin lymphoma | 17 (9.8%) |
| Acute lymphoid leukemia | 12 (6.9%) |
| Other | 11 (6.4%) |
| Myeloma | 5 (2.9%) |
| Chronic lymphocytic leukemia | 3 (1.7%) |
| **Graft type** | |
| Unmodified bone marrow or PBSC | 88 (51%) |
| T-cell depleted PBSC [3] | 72 (42%) |
| Cord blood | 13 (7.5%) |
| **Intensity of conditioning regimen** | |
| Ablative | 100 (58%) |
| Reduced intensity | 52 (30%) |
| Nonmyeloablative | 21 (12%) |

[1] Mean (SD); n (%)

[2] MDS/MPN: myelodysplastic syndromes/myeloproliferative neoplasms;

[3] PBSC: peripheral blood stem cell
